# Genomic, transcriptomic, and structural analysis of *Pseudomonas* virus PA5oct highlights the molecular complexity among Jumbo phages

**DOI:** 10.1101/406421

**Authors:** Katarzyna Danis-Wlodarczyk, Bob G. Blasdel, Ho Bin Jang, Dieter Vandenheuvel, Jean-Paul Noben, Zuzanna Drulis-Kawa, Rob Lavigne

## Abstract

*Pseudomonas* virus PA5oct has a large, linear, double-stranded DNA genome (287,182 bp) and is related to *Escherichia* phages 121Q/PBECO 4, *Klebsiella* phage vB_KleM-RaK2, *Klebsiella* phage K64-1, and *Cronobacter* phage vB_CsaM_GAP32. A protein-sharing network analysis highlights the conserved core genes within this clade. Combining genome, RNAseq and mass spectrometry analyses of its virion proteins allowed us to accurately identify genes and elucidate regulatory elements for this phage (ncRNAs, tRNAs and promoter elements). In total PA5oct encodes 462 CDS (compared to 345 *in silico* predicted genes using automated annotation pipelines), of which 25.32%, have been identified as virion-associated based on ESI-MS/MS. The RNAseq-based temporal genome organization suggests a gradual take-over by viral transcripts from 21%, 69%, and 92% at 5, 15 and 25 min after infection, respectively. Like many large phages, PA5oct is not organized into contiguous regions of temporal transcription. However, although the temporal regulation of the PA5oct genome expression reveals specific genome clusters expressed in early and late infection, many genes encoding experimentally observed structural proteins surprisingly appear to remain almost untranscribed throughout the infection cycle. Within the host, operons associated with elements of a cryptic Pf1-like prophage are upregulated, as are operons responsible for Psl exopolysaccharide (*pslE-J*) and periplasmic nitrate reductase (*napA-F*) production. The characterization described here represents a crucial step towards understanding the genomic complexity as well as molecular diversity of jumbo viruses.

## Introduction

The discovery of giant/jumbo microbial viruses infecting bacteria and protozoa has dramatically expanded the known size range of viral genomes, from around 2 kb and to over 2 Mb. Indeed, this means that jumbo viruses can be larger than the genomes of parasitic bacteria and archaea, obliterating the gulf between cells and viruses in terms of genome size and complexity (Claverie *et al.*, 2009; Koonin *et al.*, 2015). Furthermore, these viral genomes reflect an incredible viral diversity and with their large and largely uncharacterized genomes widen the already significant knowledge gap with regard to understanding the function of individual genes. Indeed, the majority of viral genes lack any nucleotide similarity to characterized or even known genes, and have aptly been described as the “Viral Dark Matter” (Hatfull, 2015).

Jumbo phages have been isolated from various environments, including water, soil, plants and animal tissues (Table S1). By our count, 108 jumbo phages have been described and sequenced to date (February 2018, Table S1), including 97 myoviruses (family *Myoviridae*), 10 siphoviruses (family *Syphoviridae*) and 1 unclassified phage. Most of these phages infect Gram-negative bacteria (95.4%), including *Synechococcus* (33 phages), *Erwinia* (20 phages), *Pseudomonas* (9 phages), *Caulobacter* (6 phages), *Vibrio* (6 phages), *Aeromonas* (5 phages), *Ralstonia* (3 phages), *Cronobacter* (2 phages), *Escherichia* (2 phages), *Klebsiella* (2 phages), *Yersinia* (1 phage), *Prochlorococcus* (1 phage), *Salmonella* (1 phage) and *Sphingomonas* (1 phage), while only 3.6% infect Gram-positive bacteria, particularly *Bacillus* strains (5 phages). Their typical tiny plaque size (< 0.5 mm on 0.7% soft agar), however, often leaves them undetected during classical phage propagation procedures, while the large size of their particles can cause them to be removed with bacteria during standard filtration steps (Serwer *et al*., 2007), thereby suggesting biases against their isolation.

Although these giant phages have proven to be excellent for structural analysis along with providing insights into infection processes (Fokine *et al.*, 2007; 2005; Wu *et al.*, 2012), the understanding of their genetic make-up and genome organization has often remained limited to the results of *in silico* predictions from genome sequencing. It is important, however, to also experimentally investigate their genome organization and transcriptional schemes as well as the mechanisms of their evolution and biology (Hendrix, 2009). RNA sequencing is proving to be an important tool in this regard, allowing both a detailed elucidation of the phage transcriptional scheme including the discovery of regulatory elements and sRNAs, but also providing clues towards the host response during phage infection and its evolutionary implications (Ceyssens *et al.*, 2014; Chevallereau *et al.*, 2016; Blasdel *et al.*, 2017b and 2018).

In this report we present a detailed analysis of the molecular aspects of the jumbo myovirus, phage PA5oct, infecting *Pseudomonas aeruginosa*. In addition, genome sequencing, temporal transcriptome and structural proteomic analyses have been combined in a comprehensive manner to experimentally elucidate the genome organization of this unique phage and the response it elicits during infection of its host. Furthermore, the evolutionary relationships between PA5oct and other (jumbo) bacteriophages are established using a protein-sharing network analysis.

## Material & Methods

### Bacteriophage propagation, purification, and morphology

In all experiments, phage PA5oct was propagated as previously described (Danis-Wlodarczyk *et al.*, 2015). Phage lysate was purified by 0.45 and 0.22 µm filtration and incubation with 10% polyethylene glycol 8000 (PEG 8000)–1 M NaCl according to standard procedures (Ceyssens *et al.*, 2006). Finally, CsCl-gradient ultracentrifugation was applied (Ceyssens *et al.*, 2008) and the resulting phage preparation dialyzed three times for 30 min against 250 volumes of phage buffer using Slide-A-Lyzer Dialysis Cassettes G2 (Thermo Fisher Scientific Inc, MA, USA). The phage titre was assessed using the double-agar layer technique (Adams, 1959) and purified samples were stored at 4°C in the dark.

### The rifampin growth assay

One-step growth curves were established according to the method of Pajunen *et al.* (2000), with modifications. An equal volume of bacterial culture at OD_600_ of 0.4 was mixed with phage suspension (10^6^ pfu/ml) to obtain a MOI of 0.01. Phages were allowed to adsorb for 8 min at 37°C, after which the mixture was diluted to 10^−4^. Triplicate samples were taken every 5 min during 1-1.5 h and titered (Danis-Wlodarczyk *et al.*, 2015). Where appropriate, rifampin (Sigma) was added to a final concentration of 400 µg/ml.

### DNA isolation and sequencing

Phage genomic DNA was prepared with the use of the modified protocol for λ DNA isolation according to Ceyssens (2009) and sequenced using the Illumina MiSeq platform available at the Nucleomics Core (VIB, Belgium), as previously described by Danis-Wlodarczyk *et al.* (2015).

### RNA extraction and sequencing

*P. aeruginosa* strain PAO1 was grown overnight in 5 ml LB medium at 37°C. Next, cells were diluted 1:100 in 50 ml fresh medium and further grown at 37°C until an OD_600_ of 0.3 was reached (∼1.2 x 10^8^ cfu/ml, early exponential phase). The culture was infected with PA5oct at a MOI of 50. To control for a synchronous infection, we ensured that less than 5% of bacterial survivors remained after five minutes post infection. Biologically independent samples were collected in triplicate at 5 minutes (early), 15 minutes (middle), and 25 minutes (late) into infection and processed as described previously (Blasdel *et al.*, 2017a; Chevallereau *et al.*, 2016).

### Genome analysis

The genome analysis was performed by integrating standard genome annotation (Danis-Wlodarczyk *et al.*, 2015; and 2016) and stranded RNA-seq reads, mapped to the phage genome as previously described (Ceyssens *et al.*, 2014; Chevallereau *et al.*, 2016). Sequencing reads were aligned separately to both the phage and host genomes using CLC Genomics Workbench v7.5.1. These alignments were then strand-independently summarized into count tables of Total Gene Reads that map to phage or host gene features respectively. Each statistical comparison presented was performed using the DESeq2 (Love et al. 2014) R/Bioconductor package to normalize host transcript populations to host transcript populations, or phage to phage, before testing for differential expression as described by Blasdel *et al.* (2018). The PA5oct genome was deposited under Genbank accession number MH121069.

### ESI-MS/MS analysis on virion particle proteins

Phage proteins were isolated from a purified phage lysate (10^9^ pfu/ml) by a single methanol/chloroform extraction (1:1:0.75, v/v/v) (Acros Organics) and precipitated by addition of an equal volume of methanol (14,000 x g, 6 min). The phage proteins were separated on a 12% SDS-PAGE ge and analyzed by ESI-MS/MS as previously described (Ceyssens *et al.*, 2014; Van den Bossche *et al.*, 2014).

### Protein family clustering and network construction and analyses dataset

To build a gene-sharing network, we retrieved 198,102 protein sequences representing the genomes of 2,010 bacterial and archaeal viruses from NCBI RefSeq (version 75) and used the network analytics tool, vConTACT (version 1.0; https://bitbucket.org/MAVERICLab/vcontact) (Bolduc *et al.*, 2017), as an app at iVirus (Bolduc *et al.*, 2017). Briefly, including protein sequences from PA5oct, a total of 198,564 sequences were subjected to all-to-all BLASTp searches, with an E-value threshold of 10^-4^, and defined as the homologous protein clusters (PCs) in the same manner as previously described (Bolduc *et al.*, 2017). Based on the number of shared PCs between the genomes, the degree of similarity was calculated as the negative logarithmic score by multiplying hypergeometric similarity P-value by the total number of pairwise comparisons. The protein-sharing relationships among the genomes with similarity values ≥ 2 (E-value ≥ 0.01) were represented. The network was visualized with Cytoscape (version 3.5.1; http://cytoscape.org/), using an edge-weighted spring embedded model, which places the genomes or fragments sharing more PCs closer to each other. Additionally, the Markov clustering (MCL) algorithm was used to group the genomes into viral cluster (VC). To maximize the cluster homogeneity, the optimal inflation factor was determined by exploring values ranging from 1.0 to 5.0 by steps of 0.2, resulting in inflation value of 1.6 with the highest homogeneity (data not shown). The taxonomic affiliation was taken from the International Committee on Taxonomy of Viruses (ICTV; http://www.ictvonline.org/virusTaxonomy) incorporated 2017 ICTV updates (Adams *et al.*, 2017; Adriaenssens *et al.*, 2017). To further uncover the substructure of VCs, the hierarchical cluster analysis was performed using unweighted pair group method with arithmetic mean (UPGMA), based on the patterns of shared gene contents (PCs) between viral genomes, where plasmids having similar gene contents patterns are placed in the same group. Further, the proportion of shared PCs between two genomes was characterized, as previously described (Bolduc *et al.*, 2017).

## Results

### Basic microbiological and genome properties of *Pseudomonas* phage PA5oct

Under standard conditions, one-step growth experiments indicate a latent period of ca. 40 min and a burst size of about 30 - 40 phage particles per infected bacterial cell (Drulis-Kawa *et al.*, 2014). When 400 µg/ml of rifampin, a host RNA polymerase (RNAP) transcription inhibitor, is added prior PA5oct infection progeny phage production is completely abolished. This suggests that PA5oct, in contrast to another giant *Pseudomonas* virus φKZ, may rely on the host RNAP for phage transcription or alternatively that an unknown but essential phage protein is also sensitive to rifampin.

Phage PA5oct has a linear, A+T-rich (33.3% GC), double-stranded DNA genome of 287,182 bp. With *in silico* predictions made with a combination of GeneMark S (Besemer *et al.*, 2001), GeneMark.hmm (Lukashin *et al.*, 1998), OrfFinder (Sayers *et al.*, 2011) and manual inspection we were able to identify 462 putative CDS that could be validated by RNA-Seq data. In contrast, the automated RAST pipeline (McNair *et al.*, 2018) was only able to identify 345 CDS (74%). Of these 345 CDS, 21 CDS (6%) could be discarded as implausible and additional 23 CDS (6%) could be annotated more strongly with an alternative start codon or frame based on RNA-Seq data. Twelve tRNAs [Met (CAT), Leu (TAA), Asn (GTT), Thr (TGT), Met (CAT), Leu (TAA), Arg (TCT), Met (CAT), Pro (TGG), Gly (TCC), Ser (TGA) and Ser (GCT)] could also be identified *in silico* and validated with RNA-Seq data.

### Temporal regulation of PA5oct genome expression

Reads originating from the phage and the host at each stage of infection were mapped to the phage and host genomes, revealing that PA5oct progressively comes to dominate host transcription. Indeed, PA5oct transcripts represented 21% of total non-rRNA transcripts within 5 minutes and eventually proceed to 69% and then 92% by middle (15 min) and late infection (25 min), respectively. The strand specificity of the results revealed that early transcription is almost exclusively performed on the Watson strand, with what can now be defined as early coding sequences, which classically is understood to be typically involved in transitions from host to phage metabolism (Fig. 2).

**Figure 2.**
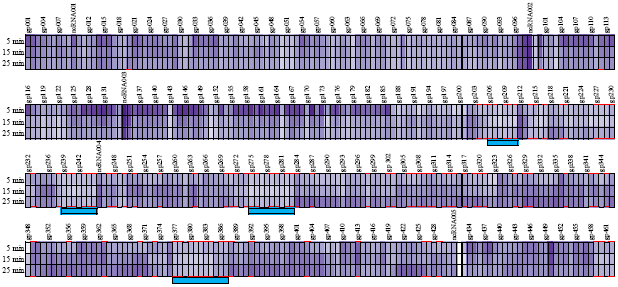
Temporal and spatial heat map summarizing PA5oct transcription by gene feature. Total gene reads aligning to each phage coding sequence and ncRNA from each of the three replicates of each condition were normalized using the DESeq normalization and their means were then divided by the Kilobasepair length of the gene feature. These expression values were then transformed on a log scale and are displayed here with lighter purple boxes representing low values and darker purple boxes representing high values. Structural genes are highlighted with red brackets and four regions of low transcription are marked with blue bars.

Most of the genes from the cluster between gp97 and gp186 (uncharacterized proteins, except gp100, gp112, and gp115, which are structural proteins), as well as gp450 (uncharacterized protein), are each found to be highly transcribed at this stage and can now be defined as early genes. Phage PA5oct appears to have a distinct middle phase of transcription captured at 15 minutes centering around the regions between gp324 - gp376 and gp388 - gp418, with gene features predicted to be responsible for structural proteins as well as DNA metabolism and replication. In the late phase of infection, gene clusters gp284 - gp376 and gp388 - gp426 are more strongly expressed compared to the early and middle phase, comprising most of the structural proteins. Additionally, five non-coding RNAs, which lack similarity to any known DNA sequence in Genbank, could be described (Table 1). Furthermore, 39 putative promoters with highly conserved, AT-rich intergenic motifs (5’-TATAATA-3’) and (5’-TTGAC-3’) were identified around transcription start sites (Fig. 1S) while 60 putative factor-independent terminators with conserved stem loops could be identified around transcription stop sites (Fig. 2S).

**Table 1.** Identified non-coding RNA species. Five non-coding RNAs have been identified based on RNA-seq data from transcripts with no plausible ORFs.

| Name | Location | Strand | Percentage of phage reads |  |  |
| --- | --- | --- | --- | --- | --- |
|  |  |  | Early | Middle | Late |
| ncRNA001 | 4113..4883 | + | 0.76% | 0.32% | 0.28% |
| ncRNA002 | 31367..31616 | + | 3.34% | 8.14% | 6.69% |
| ncRNA003 | 44676..45001 | + | 11.23% | 12.23% | 4.91% |
| ncRNA004 | 126878..127154 | + | 0.050% | 0.73% | 0.76% |
| ncRNA005 | 269893..271298 | - | 0.027% | 0.037% | 0.35% |

**Figure 1.**
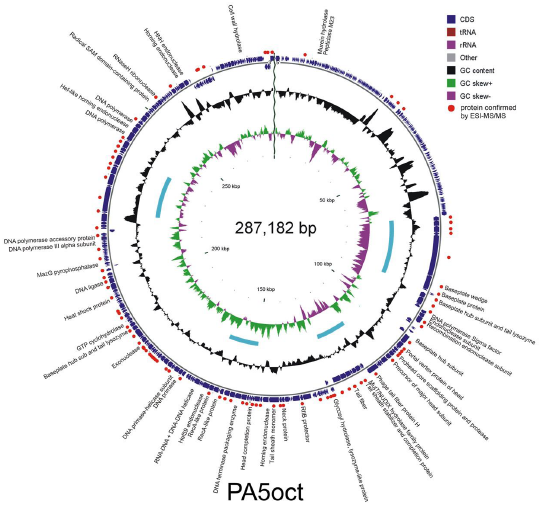
Circular representation of the PA5oct genome. Blue arrows represent predicted ORFs on the Watson and Crick strands. The inner black circle represents the GC content and the green-violet represents GC skew. Structural proteins confirmed by ESI-MS/MS analysis are marked with red dots. The blue bars represent areas with notably low transcription at each stage of infection (See Figures 2 & 3).

Remarkably, four regions, including two of the three major regions transcribed on the Crick strand (Fig. 3), were found to be only barely transcribed in spite of significant sequencing depth. Puzzlingly, each of the four regions encodes structural proteins that have been detected by ESI-MS/MS.

**Figure 3.**
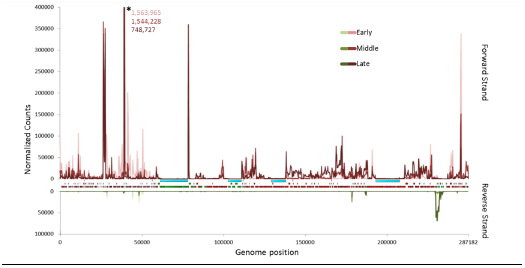
RNA-seq analysis of the PA5oct transcriptome after *P. aeruginosa* PAO1 infection. Genome-wide overview of reads mapped to the Watson (red) and Crick (green) strands of PA5oct genome for samples taken 5, 15 and 25 min post infection. Four marginally transcribed regions are highlighted with blue bars. Each of the three samples was normalized to each other by the total count of phage reads. *The asterisk represents ncRNA003, which runs off the scale of this transcriptograph.

**Figure 1S. Alignments of PA5oct promoters.** The conserved AT-rich intergenic motifs: -10 box (5’-TATAATA - 3’) and -35 box (5’ - TTGAC - 3’) are marked with pink color. The corresponding sequence logos from MEME/MAST are depicted below the alignments. ORFs in front of which promoters are located are listed on the left side, whereas the corresponding sequence locus in the phage genome is indicated on the right side.

**Figure 2S. Phage PA5oct predicted terminators with palindromes marked blue.** From the left site: list of ORFs after which terminators are located, strand, location in phage genome, motif of terminator, terminator sequence with conserved stem-loop structures marked blue, Δ*G* (the Gibbs free energy of stem-loop formation in kcal/mole).

### Transcriptome and proteome-based functional annotations

Based on bioinformatics and ESI-MS/MS analysis, the function could be predicted for 25.32 % of all proteins, including experimentally confirmed seven virion-unrelated enzymes, 13 virion-associated proteins and 73 structural gene products (Fig 3S, Table 2S). The highest amino acid similarity was found with *Cronobacter* phage vB_CsaM_GAP32, *Escherichia* phage 121Q, and *Klebsiella* phage vB_KleM_RaK2. Three capsid proteins were identified, including the portal vertex protein of the head (gp228), the prohead core scaffolding protein and protease (gp231) and the precursor to the major head subunit (gp233). An additional protein, assigned as a putative head completion protein (gp277), was annotated based on sequence similarity to the *Klebsiella* phage vB_KleM_RaK2 head completion protein. Moreover, several proteins associated with the tail apparatus were annotated, including a neck protein (gp270) and four tail proteins, a phage tail fiber protein H (gp238), a tail sheath stabilizer and completion protein (gp240), another tail fiber protein (gp243) and a tail sheath monomer (gp272) as well as three baseplate proteins, gp210, gp211 and gp214 (baseplate wedge, baseplate protein, baseplate hub subunit and tail lysozyme, respectively). Gp238, the putative phage tail fiber protein H, has a putative endo-N-acetylneuraminidase region on its C-terminus (868 – 1008 aa), suggesting a (2→8)-alpha-sialosyl linkage hydrolase function of oligo- or poly(sialic) acids, activity associated with the tail spikes and exopolysaccharide (EPS) depolymerases (Kwiatkowski *et al.*, 1982).

PA5oct also encodes several genes predicted to be associated with peptidoglycan layer degradation, including gp45, gp214, gp250, gp321 and gp447 (BLASTP, HHpred, HMMER, InterPro, Phyre2). Gp45 is a predicted murein hydrolase, peptidase M23, with a 103 amino acids long domain (30 - 133 aa). Gp214 is a putative baseplate hub subunit with tail-associated lysozyme (ESI-MS/MS hit) homologues to gp05 of *E. coli* bacteriophage T4 and which has a needle-like structure attached to the end of the tail tube serving in the injection process (Kanamaru *at al.*, 2005). Gp250 was assigned as glycosyl hydrolase, lysozyme-like protein with a 159 amino acids long domain (11 - 170 aa). Gp321 is a putative baseplate hub and tail muramoyl-pentapeptidase (ESI-MS/MS hit). Finally, gp447 is a cell wall hydrolase domain protein with an N-acetylmuramyl-L-alanine amidase SleB domain cleaving the bond between N-acetylmuramoyl and L-amino acid residues. It is probably an endolysin since it is was not recognized during ESI-MS/MS analysis.

**Figure 3S. The SDS-PAGE pattern of PA5oct structural proteome against LMW Ladder (Thermo Scientific) in the first lane.** The corresponding molecular weight is mentioned left. The numbered fractions on the right, correspond to gel slices analyzed individually by ESI-MS/MS. The proteins are mentioned in the slice in which they were most abundantly present.

**Table 2S. The ESI-MS/MS analysis of denatured phage PA5oct particles after fractionation on SDS-PAGE gel.**

### Gene-sharing relationships with other jumbo phages including PA5oct to other viruses

To represent genetic relationships between *Pseudomonas* phage PA5oct and other bacteriophages with genomes exceeding 200 kbp (Hendrix, 2009), a gene-sharing network was built. Viruses that do not show significant similarity to the 46 jumbo phages (Genbank dataset up to 2016) were excluded for clarity. The resulting network was composed of 1,355 viral genomes (nodes) belonging to the *Myoviridae, Siphoviridae, Podoviridae*, or uncharacterized phages and 37,955 relationships (edges) between them.

As shown in Fig. 4A, 22 jumbo phages were placed into the largest connected component (LCC) that predominates the double-stranded DNA phages, whereas 21 phages were placed into three isolated components. A majority (19) of the 22 jumbo phages in the LCC, which includes PA5oct, fell within a highly interconnected region (subnetwork) that is rich in the *Tevenvirinae* viruses (Upper right in Fig. 4A; Table 3S). A subsequent analysis found 5 coherent groups of viral genomes (i.e., viral clusters, VCs) embedded in this subnetwork (Fig. 4B; Table 3S), of which six jumbo phages including PA5oct and *Escherichia* phages 121Q/PBECO 4, *Klebsiella* virus RaK2, *Klebsiella* virus K64-1, and *Cronobacter* virus GAP32 were grouped into VC_77, due to the higher level of shared genes (i.e., homologous protein clusters, PCs) than the rest of network (Bolduc *et al.*, 2017). Indeed, comparison of their connectivities (i.e., the strength of connection based on the edge weight) revealed that these six viruses show stronger connections to each other but weaker connections outside the group (Fig. 4B). Notably, a further inspection of VC_77 (phage K64-1 was excluded as it has an incomplete protein annotation profiles in NCBI) uncovered its sub-structure where PA5oct shares less common PCs (∼11-15%) to the remaining members (Fig. 4C), indicating its distant relationships.

**Fig. 4.**
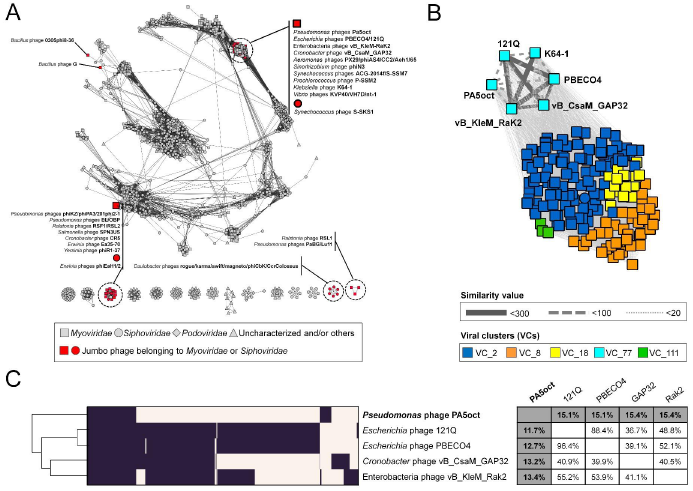
Protein-sharing network for PA5oct and jumbo phages. Protein-sharing network for PA5oct and other 46 jumbo phages. (A) A network representation was produced using the edge-weighted spring embedded layout of Cytoscape version 3.5.1. Each node is depicted as a different shape, representing bacteriophages belonging to the *Myoviridae* (rectangle), *Podoviridae* (diamond), *Siphoviridae* (circle), or uncharacterized phages (triangle). Selected jumbo phages are shown in red. Edges between two nodes indicate their statistically weighted pairwise similarities with similarity scores of ≥2. (B) An enlarged view of the subnetwork comprising PA5oct and its relatives. Edge thickness is proportional to similarity values estimated with the hypergeometric equation (Materials and methods) and viral clusters are represented in the legend boxes. (C) Module profiles showing the presence (dark) and absence (light) of homologous protein clusters (PCs) across genomes. Each row represents a virus and each column represents a PC. The genomes were hierarchically grouped based on hierarchical patterns of gene sharing (Left). A matrix showed the percentage of shared PCs between five member viruses within VC_77 (Right). From these proteome-based analyses, phage K64-1 was excluded due to its incomplete protein annotation in NCBI.

With respect to other jumbo phages, the network identifies informative connections. For example, the *φCbKvirus*, and phages RSL1, PaBG, and Lu11, which belong to VCs 87, and 127, respectively (Table 3S), form two isolated components (Fig. 4A). This discontinuous structure of two viral groups, due to their distinct gene pools, can reflect their evolutionary relationships as separate viral lineages (Gill *et al.*, 2012; Yamada *et al.*, 2010); whereas, interestingly, the inclusion of jumbo phages and their relatives having smaller genomes (<200kbp) into the same clusters, such as VCs 43 (*Bacillus* phage G), 44 (phages of the *Phikzvirus, Elvirus, Agrican357virus*, and *Rsl2virus*), and 184 (*Bacillus* phages 0305φ8-36) (Table 3S) can support the evolutionary links between jumbo phages and smaller-genome phages (Hendrix, 2009; Adriaenssens *et al.*, 2012; Jang *et al.*, 2013). Together, visualization and analysis of gene content relationships of all 46 jumbo phages as a network revealed the global distribution of genetically diverse groups of jumbo phages across 2,010 viruses, in which most of them together with smaller-genome phage(s) are placed into two major groups comprising viruses belonging to the *Tevenvirinae* as well as the *Phikzvirus*/*Elvirus*/*Agrican357virus/Rsl2virus*, respectively, and others form evolutionary distinct clades (Gill *et al.*, 2012; Yamada *et al.*, 2010) (Table 3S). In particular, despite the similarities of PA5oct to a recently emerged jumbo phages vB_CsaM_GAP32, 121Q, PBECO 4, vB_KleM-RaK2, and K64-1(Abbasifar *et al.*, 2014), follow-up analyses indicate that PA5oct appears to be more diverged with the less shared gene contents.

**Table 3S. List of jumbo phages (yellow color) and relevant phages that belong to the viral clusters (VCs) and their ICTV taxonomy.**

### Stresses uniquely imposed by PA5oct on the host as revealed by host differential expression

The conserved PAO1 host-mediated transcriptional response to phage infection present during infection by phages 14-1, PEV2, YuA, and LUZ19 (Blasdel *et al.*, 2018) also applies to the lytic infection cycle of PA5oct. However, PA5oct also appears to elicit a number of phage-specific responses on host transcription, which result in a unique differential expression pattern of specific host genes. Notably, PA5oct progressively dominates the non-ribosomal RNA environment of the cell with strong transcription, as well as the expression of an RNase-H-like protein, putatively involved in degrading of host transcripts. This shift from host transcripts to phage transcripts (Figure 5) de-enriches all host transcripts relative to the total in the cell 0.08 fold. This global depletion of host transcripts relative to the total phage/host transcript population is not represented in the Log_2_(Fold Change) values we report (Figure 5, Table 4S) as described in (Blasdel *et al.*, 2018). Briefly, this allows us to illustrate the impact of PA5oct on host transcripts relative to other host transcripts, rather than relative to the total. Remarkably, both the phage and host-mediated transcriptional stress responses abruptly cease to function around 5 minutes into infection, possibly reflecting a mechanism by which PA5oct inhibits the transcription of host genes, similar to the *Autographvirinae* (Bae *et al.*, 2013; Klimuk *et al.*, 2013), while using phage proteins to continue hijacking the host RNA polymerase.

**Figure 5.**
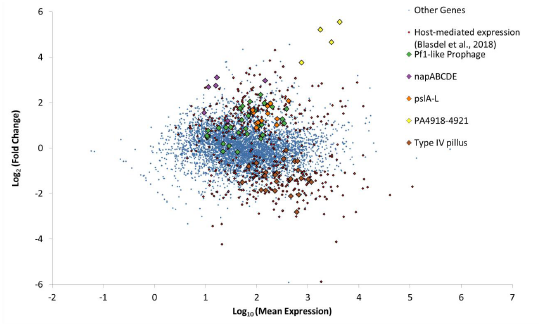
MA Plot highlighting the differential expression of host genes during the course of PA5oct infection. In addition to the host-mediated stress response to phage infection identified by Blasdel et al. (2018), we observe PA5oct-specific differential expression of host genes during PA5oct infection.

**Table 4S. The differential expression of each non-rRNA gene feature within *Pseudomonas* PAO1 genome by late PA5oct infection.**

One operon upregulated uniquely during PA5oct infection encodes the gene cluster *pslE-J,* which is involved in the production of the Psl exopolysaccharide (Jackson *et al.*, 2004). PA5oct uniquely manipulates the host into upregulating the transcription of the *napABCDEF* operon, which encodes a periplasmic nitrate reductase.

PA5oct mobilizes the transcription of genes of a potentially cryptic Pf1-like prophage encoded from ORF PA0618 to PA0642. As PA5oct infection does not upregulate the transcription of the whole prophage, and the upregulation relative to other host transcripts is not strong enough to overcome the global de-enrichment of host transcripts generally relative to phage transcripts, it is unlikely that this results in successful prophage induction, but might be related to phage-encoded superinfection exclusion mechanisms.

At the same time, a number of genes associated with the biosynthesis of the Type IV pili are downregulated relative to other host transcripts, even before considering the global replacement of host transcripts with phage transcripts. As phage PA5oct also uses the Type IV pili, as well as other factors, as receptor binding sites, this may also reflect a phage-mediated mechanism. Indeed, by reducing the number of phage binding sites, PA5oct would be able to prevent the loss of “sister” phages by phage adsorption inhibition of already phage-infected bacteria (secondary adsorption) (Abedon, 2017).

## Discussion & conclusions

An unique *Pseudomonas* phage PA5oct is a member of an emerging clade of giant phages. It appears to represent a new genus based on genome organization, phylogenetic clustering and the existing parameters for taxonomic guidelines currently being applied by the ICTV bacterial virus subcommittee (A. Kropinski, personal communication). This clade includes *E. coli* 121Q, PBECO4, *Klebsiella* phage RaK2 and *Cronobacter* phage GAP32. The representatives of this group belong to the *Myoviridae* family, have very large heads, relatively short tails, and limited genomic relationships with other phages (Abbasifar *et al.*, 2014).

### Comparative genome analysis and protein-sharing network

To examine the genetic relationships between PA5oct and other jumbo bacteriophages as well as *Myoviridae, Siphoviridae, Podoviridae* viruses, a protein-sharing network was constructed. The PA5oct, *Escherichia* phages 121Q/PBECO4, *Klebsiella* phage vB_KleM-RaK2, *Klebsiella* phage K64-1, and *Cronobacter* phage vB_CsaM_GAP32 presented closer relationships based on shared conserved core genes. Furthermore, PA5oct, PBECO4, GAP32, T5, FelixO1 appears to be distantly diverged members of *Tevenvirinae*. The T5 and FelixO1 phages appear also to act as bridge for T4-related components due to their links to other phage groups. Unlike the marker genes of bacteria (i.e., 16S rRNA genes), which can be used for their taxonomic classification, viruses lack universal genes. Thus, traditional single-gene-based phylogenies will always be limited to within only relatively closely related subsets of total viruses. Protein-sharing networks, as presented here, introduce an alternative approach, which provides a more global view which is also consistent with current taxonomic groupings. Using this approach, the possible genetic relationships for not only PA5oct but all giant phages among whole phage population could be observed.

### Genome properties

Phage PA5oct DNA sequencing revealed a linear double-stranded DNA genome, which is the ninth largest sequenced myovirus and third largest among *Pseudomonas* jumbo phages, that also includes 201φ2-1 (316.674 kbp, Thomas *et al.*, 2008), ΦPA3 (309.208 kbp, Monson *et al.*, 2011), OBP (284.757 kbp, Cornelissen *et al.*, 2012), Lu11 (280.538 kbp, Adriaenssens *et al.*, 2012), φKZ (280.334 kbp, Mesyanzhinov *et al.*, 2002), KTN4 (279.593 kbp, Danis-Wlodarczyk *et al.*, 2016), PA7 (266.743 kbp, Kwan *et al.*, 2006), PaBG (258.139 kbp, Sykilinda *et al.*, 2014) and EL (211 kbp, Krylov *et al.*, 2003).

Importantly, to date, the largest myovirus known is *Bacillus megaterium* phage G (497,513 bp genome, Table 1S). However, some giant phages, especially those reported prior to the 1990s, were identified only by electron microscopy (e.g. *Gluconobacter* phage GW6210) (Abbasifar *et al.*, 2014).

Unfortunately, a putative function can be predicted for only a small number of phage PA5oct proteins, which similarly has been the case for other jumbo phage genomes. A large portion of the predicted CDS are unique, without any sequence similarity in the currently available databases. For this reason the functional annotation of giant phage genomes is of utmost importance to elucidate basic phage biology properties as well as phage application safety (Yuan *et al.*, 2017). The whole genome analysis of transcription using RNA seq is a powerful way to elucidate differential expression of gene features across different conditions. By experimentally defining the timing and expression levels of transcripts in both phage and host, directional RNA seq has the ability to discover novel coding sequences, particularly for non-coding RNAs and small phage peptides falling below gene prediction thresholds (Ceyssens *et al.*, 2014). In this manner, we can refine annotations of existing coding sequences, predicted strictly *in silico* as based on the presence of open reading frames and often distant orthology to other often hypothetical features. In the case of PA5oct, RNA sequencing idenified 122 additional ORFs (+26%) and confirmed the presence of previously predicted genes. In total, 462 ORF’s were identified. The highest amino acids similarity was found with *Cronobacter* phage vB_CsaM_GAP32, *Escherichia* phage 121Q, and *Klebsiella* phage vB_KleM_RaK2.

Giant phages also have a more complex structure of their virions in comparison to smaller phages. ESI-MS/MS allowed us to identify 87 structural proteins of PA5oct, including four structural head related proteins, one neck protein and 4 tail related proteins. This is a similar result to that seen with giant *Pseudomonas* 201φ2-1 (89 structural proteins, Thomas *et al.*, 2010). However, other jumbo phages can possess a much smaller number of structural proteins, such as *Pseudomonas* phage ΦKZ (62), *Aeromonas* phage phiAS5 (26) or *Ralstonia* phage phiRSL1 (25) (Lecoutere *et al.*, 2009; Yamada *et al.*, 2010; Kim *et al.*, 2012). Furthermore, the capsid of jumbo phages can have very complex structures, as we could observe in the case of phage ΦKZ (30 head structural proteins), where in the middle it has an “inner body”, a spool-like protein structure that plays an important role in DNA packaging and genome ejection during, respectively, phage virion assembly and subsequent virion adsorption (Wu *et al.*, 2012).

The large capsids and consequently large genomes allow jumbo phages to possess many accessory genes comparing to small phages (Ceyssens *et al.*, 2014; Yuan & Gao, 2017; Mesyanzhinov *et al.*, 2002; Hertveldt *et al.*, 2005; Kiljunen *et al.*, 2005; Thomas et al., 2007; Gill *et al.*, 2012). For example, PA5oct encodes five different genes encoding peptidoglycan-degrading enzymes and 12 tRNA genes to increase the translation efficiency (Yuan & Gao, 2016). However, other jumbo phages contain many more tRNAs genes, such as *Vibrio* phage φ-pp2 and KVP40 (30 tRNA); *Caulobacter* phage CcrColossus and *Vibrio* phage nt-1 (28 tRNA); *Aeromonas* phage Aeh1, *Caulobacter* phage Swift and Magneto (27 tRNA); *Cronobacter* phage GAP32, *Caulobacter* phage Karma and φCbK (26 tRNA); *Aeromonas* phage PX29 (25 codons) (Table 1S).

### Hijacking the transcriptional environment of the host

The strategy of phage transcriptional mechanisms depends primarily on the presence or absence of phage RNA polymerase (Yang *et al.*, 2014). After careful analysis of PA5oct genes, no coding sequences that could be predicted *in silico* to have RNAP activity could be identified. This together with the absence of phage production in the presence of rifampicin suggest that this phage may depend entirely on host transcriptional mechanisms, even though such dependence likely represents both an under-identified and very common feature among jumbo phages. The main role in phage genes transcription plays large DNA-dependent RNAP of bacterial host, a 400-kDa protein complex composed of five subunits (α_2_, β, β’,ω), that transcribes mRNA following binding to DNA template (Murakami, 2015; Tagami *et al.*, 2014, Davis *et al.*, 2017). This process is often supported by phage-encoded σ factor forming a holoenzyme with RNAP (α_2_, β, β’,ω, σ) (Williams *et al.*, 1987; Pavlova *et al.*, 2012). However, phages can implement other transcriptional strategies, e.g., viral gene transcription may rely on both the host RNAP along with a single-subunit phage RNAP (*Enterobacteria* phage T7, T3, *Pseudomonas* phage phiKMV and *Xanthomonas oryzae* phage Xp10). The single-subunit RNAP is responsible for transcription of phage genes in the middle and late stage of infection, whereas early genes are transcribed by the host RNAP (Savalia *et al.*, 2010; Semenova *et al.*, 2005, Yang *et al.*, 2014). A similar situation is observed in the case of *Enterobacteria* phage N4, where early and middle stage of transcription depends on two phage-encoded RNAPs and the late genes are transcribed by host RNAP (Haynes *et al.*, 1985; Willis *et al.*, 2002). The giant *Pseudomonas* phage ΦKZ, can infect its host independently of the host RNAP using a virion-associated phage RNAP active in early transcription, and a non-virion-associated phage RNAP, important for late and possibly middle transcripts (Ceyssens *et al.*, 2014; Yakunina *et al.*, 2015). Our RNA seq data analysis also shows that phage PA5oct expresses a host RNase-H-like protein, that is putatively involved in degradation of host transcripts.

### Phage transcription patterns

Generally, dsDNA tailed phages follow different temporal expression stages during lytic cycle. In the early phase, genes responsible for the host defense disarmament and host metabolism conversion towards viral protein production are expressed. This group of genes are the most diverse and often do not have assigned functions. Next, transcription focuses on genes involved in DNA replication and nucleotide biosynthesis. In the late stage, genes associated with DNA packing, phage morphogenesis and lysis of the host cell are overexpressed (Lavysh *et al.*, 2017). However, similar to giant *Pseudomonas* phage ΦKZ (Ceyssens *et al.*, 2014), giant *Yersinia* phage ΦR1-37 (Leskinen *et al.*, 2016) and giant *Bacillus* phage AR9 (Lavysh *et al.*, 2017), but unlike most smaller tailed phages, PA5oct is not organized into contiguous regions of temporal transcription. Indeed, regions of early, middle, and late expression are scattered throughout the genomes as increasingly appears to be characteristic of giant phages. We also found four regions with puzzlingly low levels of transcription that encode for structural genes correlated to baseplate and tail (gp206-gp211: baseplate proteins, gp239-gp245: tail sheath and tail fiber, gp275-gp283 and gp377-gp387) that were confirmed to be translated and incorporated into phage particles by ESI-MS/MS. Notably, even the inadequately low levels of transcription detected for these regions do not appear to be rationally temporally regulated as they are most strongly present at 5 and 25 minutes, but almost disappear at 15 minutes into infection. They are also not associated with an evolutionary history which is distinct relative to the rest of the phage genome, ruling out the possibility that they represent recently acquired phage mosaic tiles.

Within the infection, PA5oct progressively dominates host transcription in a constant rate. After 5 min, PA5oct transcripts represented 21% of total non-rRNA transcripts, eventually proceed to 69% then 92% by middle and late infection, respectively. This possibly reflects a globally accelerated degradation of RNA in the cell similar to what was described by Chevallereau *et al*, (2016).

### Host stress response during PA5oct phage infection

Phages have a substantial impact on bacterial cellular systems, including transport/export, energy production/conversion, ribosomal proteins, cell wall modification, conversion of ribonucleotides to deoxyribonucleotides, cold shock and osmotic stress response (Fallico *et al.*, 2011; Ainsworth *et al.*, 2013; Ravantti *et al.*, 2008; Leskinen *et al.*, 2016). In this study, we observed that PA5oct upregulates the operon encoding *psIE-J*, responsible for production of Psl exopolysaccharide. Psl is essential for initiating and maintaining biofilm architecture in both mucoid and non-mucoid strains. It is responsible for the formation of a fabric-like matrix that holds cells closely together, thereby impeding migration within the biofilm (Ma *et al.*, 2009). This may lead to increased slime production, reducing the relative abundance of LPS phage receptors, inhibiting adhesion by sister phages.

Also the *napABCDEF* operon, involved in production of a periplasmic nitrate reductase, is upregulated during PA5oct infection. The periplasmic nitrate reductase is required for anaerobic growth in cystic fibrosis (CF) sputum *in vitro.* In CF environment, *P. aeruginosa* prefers anaerobic growth that significantly enhances its biofilm formation and antibiotic resistance (Yoon *et al.*, 2002). Under these anaerobic conditions, *P. aeruginosa* modifies the structure of its LPS from a highly electronegative surface to a neutral surface (Sabra *et al.*, 2003, Palmer *et al.*, 2007). This suggests that phage PA5oct may influence *Pseudomonas* biofilm formation. While the purpose and mechanistic origin of this upregulation is unclear, it would seem to suggest that PA5oct may be well adapted to the environment of human-associated *Pseudomonas* biofilms.

Since PA5oct downregulates biosynthesis of IV Type pili and indirectly modifies bacterial LPS, the signaling networks might also influence the overproduction of nitrate reductase. As it was proved in our back to back paper IV Type pili and LPS structures serve as phage PA5oct receptors [Olszak *et al.*, bioRxiv https://doi.org/10.1101/405027]. Receptor downregulation possibly functions as a tactic to reduce losses of “sister” phages to adsorption (i.e., secondary adsorption) of already phage-infected bacteria (Abedon, 2017). By not suicidally adsorbing already phage-infected bacteria, free phages released by nearby phage-infected bacteria thereby are able to instead target uninfected bacterial cells embedded deeper in the biofilm (that is, beyond more surface-located infected bacteria), to acquire biofilm dispersal cells, or to simply diffuse away from the biofilm towards seeking more distantly located bacteria to infect (Høyland-Kroghsbo *et al.*, 2013; Abedon, 2017). Such interference with phage secondary adsorption would occur independently from mechanisms of superinfection exclusion or superinfection immunity, as have been found to be displayed by phage infected bacteria in other systems, and both of which act *following* secondary adsorption (Abedon, 2015, 2017). Thus, rather than strictly a mechanism of prevention of superinfection, and thereby in some manner protecting the phage-infected bacterium, infected bacteria by interfering with secondary adsorption also, at least in principle, may protect – from suicidally secondarily adsorbing – clonally related “sister” PA5oct virions released nearby within the same biofilm.

We also observed that PA5oct stimulates transcription of genes associated with prophage elements within its host. Prophage transcription appears to be ubiquitous during *Pseudomonas* phage infection, even in heavily domesticated lab strains, suggesting that a superinfecting lytic phage may need to compete with these other viruses for metabolic control of the cell. This was also previously observed in case of other phages, such as LUZ19, PEV2, ΦKZ, PakP3, and PakP4 (Blasdel *et al.*, 2018).

### RNAseq as a useful tool for genome annotation

Whole genome analysis of transcription using RNA seq is a powerful way to elucidate differential expression of gene features across different conditions and otherwise revolutionized genome annotation. With directional RNA seq, as demonstrated here, starting with strictly *in silico* predictions, i.e., as based on the presence of open reading frames and often distant orthology to other often hypothetical features, we can substantially refine annotations of existing protein coding sequences and sRNAs.

## Funding

This study was supported by research grant no. 2012/04/M/NZ6/00335 of the National Science Centre, Poland. KDW was co-financed by the European Union as part of the European Social Fund and a postdoctoral fellowship at KU Leuven, Belgium. RL is a member of the “PhageBiotics” research community, supported by the FWO Vlaanderen. ZDK was a members of the EU COST BM1003 Action in Microbial cell surface determinants of virulence (http://www.cost.eu/COST_Actions/bmbs/BM1003). JPN is supported by a grant from the Hercules Foundation (project R-3986).

### Acknowledgements

We are grateful to Stephen T Abedon for critical comments and helpful suggestions on the manuscript.

KDW performed one-step growth experiments, DNA isolation, genome annotation and analysis of ESI-MS/MS results. HBJ generated protein family clustering, network construction and analyzed the dataset. DV helped with promoter identification. JPN performed ESI-MS/MS analysis. KDW and BB prepared samples for RNA seq and analyzed results. KDW, HBJ, BB, RL and ZDK designed experiments and wrote the manuscript.

